# *Helicobacter pylori* chronic infection selects for effective colonizers of metaplastic glands

**DOI:** 10.1101/2022.06.13.496036

**Authors:** VP O’Brien, LK Jackson, JP Frick, AE Rodriguez Martinez, DS Jones, CD Johnston, NR Salama

**Affiliations:** Human Biology Division, Fred Hutchinson Cancer Center, Seattle, WA, USA; Molecular and Cellular Biology Graduate Program, University of Washington, Seattle, WA, USA; Department of Microbiology, University of Washington, Seattle, WA, USA; Vaccine and Infectious Disease Division, Fred Hutchinson Cancer Center, Seattle, WA, USA

## Abstract

Chronic gastric infection with *Helicobacter pylori* can lead to progressive tissue changes that culminate in cancer, but how *H. pylori* adapts to the changing tissue environment during disease development is not fully understood. In a transgenic mouse gastric metaplasia model, we found that strains from unrelated individuals differed in their ability to infect the stomach, to colonize metaplastic glands, and to induce proliferation and alter the expression of metaplasia-associated proteins. *H. pylori* isolates from different stages of disease from a single individual had differential ability to colonize healthy and metaplastic gastric glands. Exposure to the metaplastic environment selected for high gastric colonization by one of these strains. Complete genome sequencing revealed a unique alteration in the frequency of a variant allele of the putative adhesin *sabB*, arising from a recombination event with the related sialic acid binding adhesin (SabA) gene. Mutation of *sabB* strongly reduced adherence to both normal and metaplastic gastric tissue in multiple strain backgrounds and highly attenuated stomach colonization. Thus, the changing gastric environment during disease development promotes bacterial adhesin gene variation associated with enhanced gastric colonization.

**Importance:** Chronic infection with *Helicobacter pylori* is the primary risk factor for developing stomach cancer. As disease progresses *H. pylori* must adapt to a changing host tissue environment that includes induction of new cell fates in the cells that line the stomach. We tested representative *H. pylori* isolates collected from the same patient during early and later stages of disease in a mouse model where we can rapidly induce disease-associated tissue changes. Only the later-stage *H. pylori* strains could robustly colonize the diseased stomach environment. We also found that the ability to colonize the diseased stomach was associated with genetic variation in a putative cell surface adhesin gene called *sabB*. Additional experiments revealed that SabB promotes binding to stomach tissue and is critical for stomach colonization by the late-stage strains. Thus, *H. pylori* diversifies its genome during disease progression and these genomic changes highlight critical factors for bacterial persistence.

## Introduction

*H. pylori* stomach infection is the leading risk factor for the development of gastric malignancies, with an estimated 75% of gastric cancer cases attributed to active infection ^1^. Although the mechanisms by which *H. pylori* induces gastric cancer are incompletely understood, the tissue changes preceding cancer development have been well described ^2^. All infections lead to chronic inflammation of the stomach lining (gastritis). A fraction of these cases progress to atrophic gastritis (loss of acid secreting parietal cells), SPEM (spasmolytic polypeptide expressing metaplasia) and/or IM (intestinal metaplasia), which then can progress to dysplasia and finally gastric cancer ^3, 4^. Eradication of *H. pylori* prior to the development of metaplasia reduces the risk of gastric cancer development ^5^, which supports a model in which *H. pylori* promotes early tissue alterations leading to carcinogenesis.

The interactions between host and pathogen are dynamic throughout infection as acid homeostasis, nutrient availability, and immune responses fluctuate ^6^. Infections are typically established in the stomach antrum, where pH is closer to neutral. However, loss of acid secreting parietal cells due to atrophy of the stomach corpus glands promotes expansion of *H. pylori’s* niche from the antrum to the corpus in both human and animal models ^7, 8^. Within these different topographic regions, *H. pylori* can be found in the protective mucus layer above the epithelium and within glands ^9–11^. At the epithelial interface, *H. pylori* secretes effectors directly into host cells as well as into the surrounding environment, resulting in a robust inflammatory response ^12^. It is thought that by-products of inflammation generated from infection lead to accumulation of mutations in gastric epithelial cells, which are sufficient to activate oncogenic pathways in some individuals ^13^.

We recently showed that *H. pylori* presence alters the trajectory of SPEM, IM, and dysplasia ^14^, in addition to its previously described role in driving inflammation associated with oncogenic mutations. These studies suggest that *H. pylori* modulates pre-malignant tissue changes, which in turn can impose selective pressures on the infecting bacterial populations. *H. pylori* persistence in response to tissue remodeling in the gastric environment is facilitated by genomic diversification through mutation as well as inter and intra genomic recombination ^15^. Comparisons of paired *H. pylori* isolates collected longitudinally from the same individual document extensive genetic diversity, suggesting adaptation to distinct niches within a single host ^16–18^. However, few studies have characterized how *H. pylori* specifically interacts with, and may genetically adapt to, the metaplastic tissue environment. Here we utilized a mouse model of KRAS-driven metaplasia to explore bacterial genotypes that promote colonization of the metaplastic stomach.

## Results

### The model H. pylori strain PMSS1 robustly colonizes the gastric corpus during gastric preneoplastic progression

We previously used *Mist1-Kras* mice to explore how *H. pylori* infection could impact metaplasia development in the stomach corpus ^14^. In these mice, administration of tamoxifen (TMX) induces expression of an active *Kras* allele (G12D) in chief cells, leading to rapid onset of spasmolytic polypeptide-expressing metaplasia (SPEM) in the corpus within four weeks, which progresses to intestinal metaplasia (IM) and mild dysplasia by 12 weeks ^19^. In the present study we either induced active KRAS (KRAS+), or sham-induced mice (KRAS-), and then performed acute *H. pylori* infections to assess how *H. pylori* interacted with metaplastic vs. healthy glands (**Figure 1A**). We first tested *H. pylori* strain PMSS1, which robustly colonizes the healthy mouse stomach ^8, 20^. Mice were infected for one week during SPEM (four weeks after KRAS induction), SPEM with intestinalizing characteristics (SPEM-IC, eight weeks after KRAS induction), or IM (12 weeks after KRAS induction), with sham-induced mice as a control at each time point (**Figure 1B**). PMSS1 robustly colonized the KRAS-mice at each time point, in keeping with its previously established ability to colonize wild-type mice. This strain also robustly colonized KRAS+ mice at each time point, demonstrating that it could survive in the metaplastic stomach. Interestingly, at 12 weeks, some KRAS+ mice had bacterial loads one to two logs higher than sham induced mice. We hypothesized that this could result from *H. pylori* expansion from the antrum into corpus metaplastic glands.

**Figure 1.**
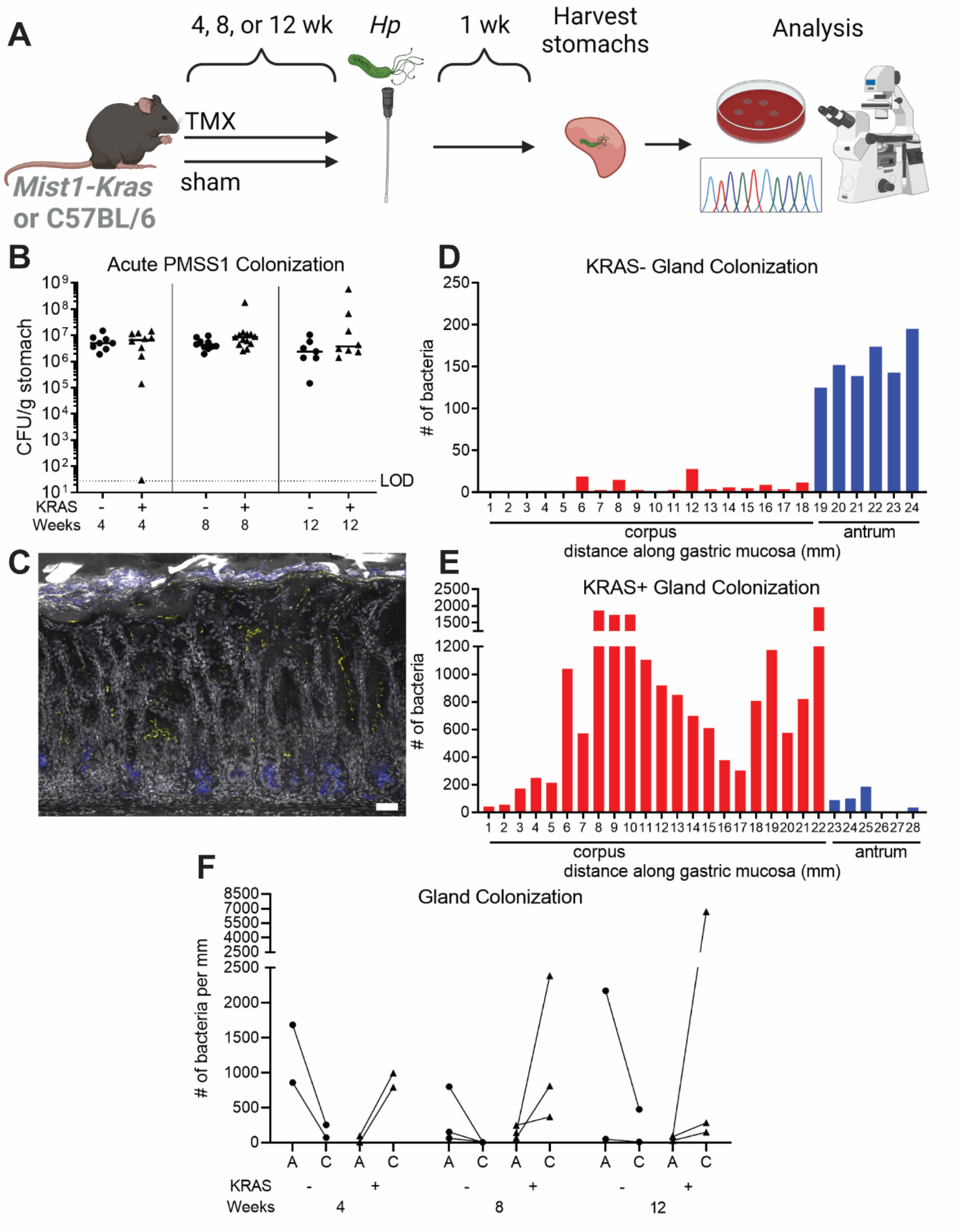
*H. pylori* colonizes the stomach corpus during gastric pre-neoplastic progression. (**A**) *Mist1-Kras* mice were given tamoxifen (TMX) to induce active KRAS in the chief cells, which leads to metaplasia development, or were sham-induced with vehicle (corn oil). After the onset of SPEM (4 weeks post-TMX), SPEM with intestinalizing characteristics (8 weeks), or intestinal metaplasia (12 weeks), mice were infected with *Helicobacter pylori* for one week to assess acute *H. pylori*-tissue interactions. Illustration created with BioRender.com. (**B**) At the indicated weeks after TMX treatment (+) or sham injection (-), mice were infected with PMSS1 for one week. Bacterial titers of individual mice are shown and a bar indicates the median. Data are from N=2 independent experiments with n=7-13 mice per group. CFU, colony-forming units; LOD, limit of detection. (**C-E**) Thick stomach sections (200 µm) were stained for *H. pylori* and imaged via confocal microscopy. Z-stacks were collected along the entire length of the stomach and volumetric analysis was used to enumerate bacteria based on fluorescent voxels. (**C**) Representative maximum intensity projection of a stomach section from the corpus of a mouse infected with *H. pylori* strain PMSS1 for one week, 12 weeks after KRAS induction. Grey, DAPI; yellow, *Hp*; blue, GS-II (metaplasia marker). Scale bar, 50 μm. (**D-E**) Shown are representative examples of gland analysis for PMSS1 in KRAS- (**D**) and KRAS+ (**E**) mice infected eight weeks after induction or sham induction. The graphs show the number of bacteria within glands as detected by fluorescent voxels, along the length of the stomach in millimeters (mm). Red bars indicate the corpus and blue bars indicate the antrum. (**F**) Mice were infected at the indicated time points, four, eight or 12 weeks after active KRAS induction or sham induction. Shown is the number of bacteria in glands per millimeter of antrum (‘A’) and corpus (‘C’) of each mouse after one week.

In humans, *H. pylori* infection initially localizes to the antrum, which lacks acid-producing parietal cells, and then gradually can spread to the corpus (main body of the stomach) over a period of years to decades ^21^. A similar phenomenon was observed in mice: PMSS1 localized to the antrum after one week of infection in wild-type C57BL/6 mice and then spread to the corpus within one month ^8^. To visualize bacteria within glands, we used immunofluorescence microscopy to detect PMSS1 in thick sections (200 µm) of formalin-fixed tissue (**Figure 1C, Supplemental Movie 1**). We previously used immunofluorescence microscopy to detect and quantify *H. pylori* within gastric glands based on fluorescent voxels ^8^. Here we used this method to compare antral and corpus gland occupation between KRAS- and KRAS+ mice infected with PMSS1. We examined stomach sections from two to three mice per time point. At each time point, more *H. pylori* cells were present within the glands of KRAS+ mice than KRAS-mice (**Figure 1D-F**), suggesting that the metaplastic changes to the gland architecture and microenvironment may favor *H. pylori* gland colonization. Consistent with our hypothesis, far more bacteria were detected in corpus glands than antral glands of KRAS+ mice at each time point (**Figure 1F**), whereas KRAS-mice maintained the expected antral predominance. Thus, *H. pylori* strain PMSS1 can robustly colonize metaplastic corpus glands.

### Genetically diverse H. pylori strains colonize the metaplastic stomach environment

We tested a panel of *H. pylori* isolates to look for strains that might differ in their ability to colonize the metaplastic vs. healthy stomach (**Figure 2A**). NSH57 is a mouse-adapted version of the well-studied clinical isolate G27 but is a relatively poor colonizer of mice ^22, 23^. This strain colonized 0/5 KRAS-mice and 1/9 KRAS+ mice in our study. Strain 7.13 is a gerbil-adapted carcinogenic *H. pylori* strain derived from the duodenal ulcer strain B128 ^24^. Titers of 7.13 did not differ between KRAS- and KRAS+ mice. However, only 4/7 KRAS-mice and 5/8 KRAS+ mice were colonized, and median titers were three to four logs lower than titers of PMSS1, demonstrating that 7.13 is overall not a robust colonizer of mice, regardless of gastric metaplasia status. Thus, induction of metaplasia does not enhance stomach colonization by these strains.

**Figure 2.**
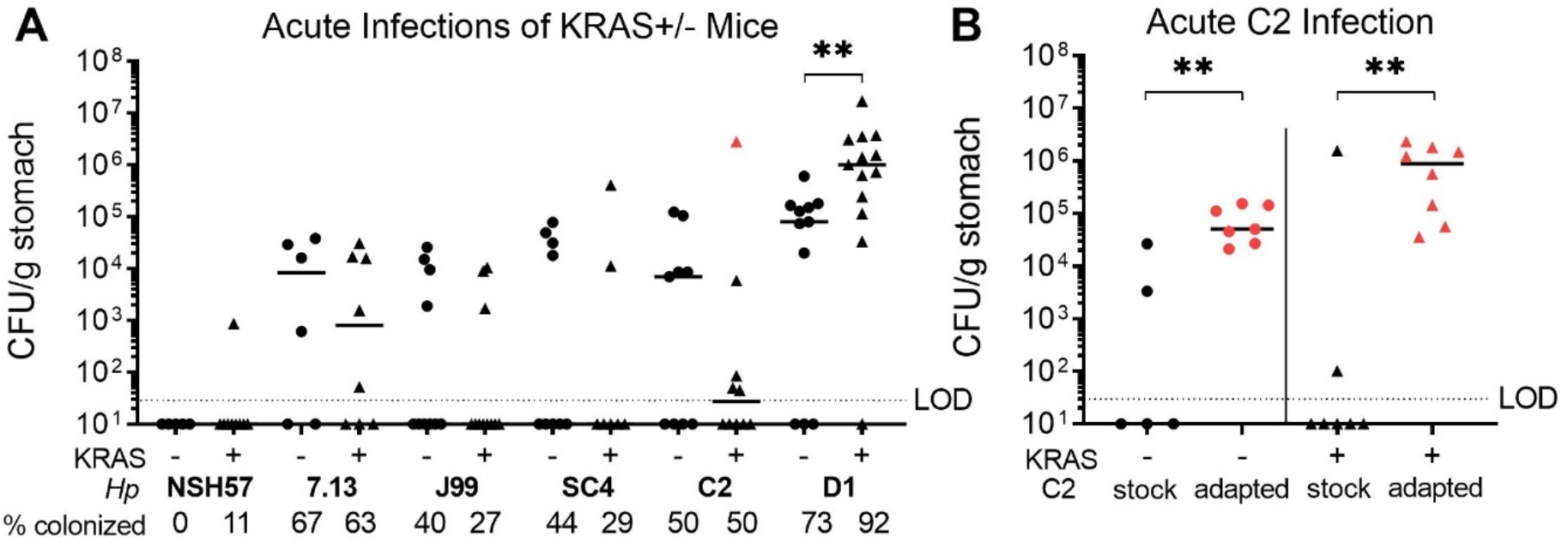
The recent isolate C2 can adapt to better colonize the mouse stomach. *Mist1-Kras* mice were sham-induced (-) or induced with TMX (+). After eight weeks, mice were infected with *H. pylori* for one week. (**A**) Shown are stomach titers after one week of infection. The percentage of total mice colonized above limit of detection (10^1^ CFU/g of stomach tissue) is listed below each isolate name. The mouse from which the C2 “adapted” strain was isolated is indicated in red and used in additional experiments (**B**) in comparison to infection with the same C2 stock culture (“stock”). Data points represent actual values from each individual mouse, bars represent the median, and mice with no detectable CFU are plotted below the limit of detection (LOD). Data are from N=2-3 independent experiments. **, *P* < 0.01, Mann-Whitney U test.

We next tested infection with representative isolates from 4 distinct genetic subgroups within the J99 strain collection of *H. pylori* isolates taken from a single human host ^25, 26^. Based on shared genetic variation, the J99 isolates cluster in four distinct subgroups ^27^. Subgroups 1A and 1B represent single colony isolates collected from an antral biopsy obtained in 1994, when the patient first presented with a duodenal ulcer. The patient refused to take the prescribed antibiotics to eradicate the infection. Subgroups 2A and 2B single colony isolates come from biopsies taken six years later from the antrum, corpus and duodenum, when the patient progressed to atrophic gastritis (a precursor to metaplasia) with loss of acid secretion ^28^. The strains J99, SC4, D1 and C2 are representative of subgroups 1A, 1B, 2A and 2B, respectively.

The J99 strain was a relatively poor colonizer of mice, with titers recovered in only 4/10 KRAS-mice and 3/11 KRAS+ mice (**Figure 2A**). The other strain from 1994, SC4, was likewise recovered from only 4/9 KRAS-mice and 2/7 KRAS+ mice. Strain D1 was recovered from 8/11 KRAS-mice and 12/13 KRAS+ mice, demonstrating that it could robustly colonize mice regardless of metaplasia status. D1 titers were about one log higher in KRAS+ mice than in KRAS-mice (*P* < 0.01, Mann-Whitney U test), suggesting that this strain preferentially colonized the metaplastic stomach. The other strain tested from the later biopsies, C2, was recovered from 5/10 KRAS-mice and 5/10 KRAS+ mice. We observed that the C2 strain was the most variable in overall bacterial burdens, with titers ranging from undetectable to greater than 10^6^ CFU/g stomach. We thus hypothesized that C2 might be poised to adapt to the metaplastic stomach environment.

To test this hypothesis, mice were infected with the C2 “stock” strain (as in **Figure 2A**) or a C2 “adapted” strain recovered from a KRAS+ mouse, shown in red in **Figure 2A**. Strikingly, titers of the C2 adapted strain were significantly higher than the C2 stock strain, regardless of KRAS expression (**Figure 2B**). The C2 stock strain was recovered from 2/5 KRAS-mice and 2/7 KRAS+ mice, whereas the adapted strain was recovered from all mice tested. Thus, C2 appears poised to adapt *in vivo* to efficiently colonize the mouse stomach. Titers of the adapted C2 strain were higher in KRAS+ mice than KRAS-mice (median CFU 5×10^4^ vs. 9×10^5^, respectively, **Figure 2B**), suggesting that this adaptation may even favor the metaplastic environment.

### H. pylori isolates vary in their ability to infect metaplastic glands

Because PMSS1 exhibited robust colonization of corpus glands in KRAS+ mice (**Figure 1F**), we tested whether other *H. pylori* isolates could also colonize KRAS+ corpus glands. We stained thin sections (4 µm) with an anti-*H. pylori* antibody (**Figure 3A**) and semi-quantitatively scored the extent of bacteria in glands in a blinded fashion, where 0 indicates no bacteria detected in corpus glands, 1 indicates few bacteria detected in the corpus (0-5 per field of view), and 2 indicates moderate to abundant bacteria detected in the corpus (>5 per field of view). Only mice with detectable CFU were included in these experiments. As expected, based on our analysis of thick sections (**Figure 1E-F**), PMSS1 robustly colonized KRAS+ corpus glands (**Figure 3B and S1**). Few bacteria of any strain tested were seen in the antral glands (not shown).

**Figure 3.**
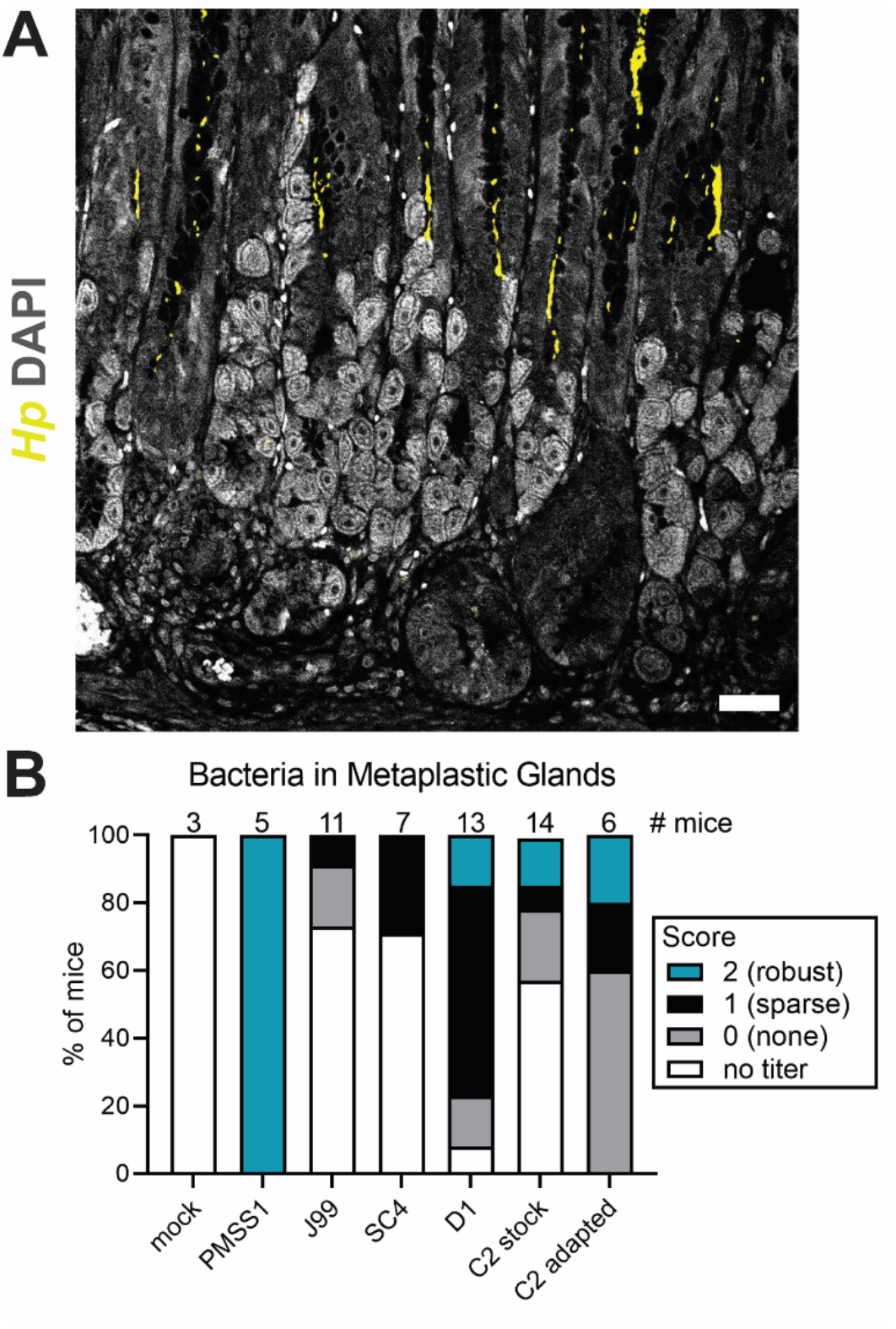
*H. pylori* strains can be detected in corpus glands at different densities. Thin stomach sections (4 µm) were stained for *H. pylori* and the corpus was assessed for bacterial colonization. (**A**) Shown is a representative image of a mouse infected with PMSS1 with a score of 2 (robust gland colonization). Yellow, *Hp*; grey, DAPI; scale bars, 50 µm. (**B**) Shown is the corpus gland colonization score for each *H. pylori* strain tested. The number of mice is given at the top of the graph. Blue indicates robust gland colonization, black indicates sparse gland colonization, and grey indicates no observable gland colonization. White indicates mice with undetectable *Hp* loads by gastric culture, which were not included in staining experiments.

Most KRAS+ mice infected with J99 and SC4 had undetectable bacterial loads (**Figure 2A**). In the few mice with detectable titers, few or no bacteria were observed within the glands (**Figure 3B**). Thus, J99 and SC4 are poor colonizers of mice and poor colonizers of metaplastic glands. Interestingly, the D1 strain fared much better: only one mouse of 13 had an undetectable bacterial titer, and bacteria were observed in KRAS+ corpus glands in 10/13 mice (**Figure S1**), while 2/13 mice had titers but did not have detectable bacteria in corpus glands. Once again, the C2 stock strain gave rise to the most variable phenotypes. Eight of 14 mice had no titers and were excluded from the analysis, but of the remaining six, three had no bacteria observable in glands, one had sparse bacteria and two had abundant bacteria. Interestingly, this pattern was similar for the C2 adapted strain. All eight mice had detectable titers (**Figure 2B**). We randomly chose five mice for gland analysis and observed that three had no bacteria in glands whereas one had sparse bacteria and one had abundant bacteria. Thus, the C2 adapted strain infects mice better than the C2 stock strain does, but in mice with detectable titers, both strains colonize metaplastic glands to a similar extent.

### H. pylori strains differentially impact metaplasia development

We previously used *Mist1-Kras* mice to determine whether chronic infection with *H. pylori* strain PMSS1 impacted metaplasia development in KRAS+ mice, compared to mock-infected mice ^14^. In these experiments, mice are infected or mock-infected prior to tamoxifen administration to induce active KRAS. We found that at six weeks, the expression of SPEM markers and the cell proliferation marker KI-67 differed between PMSS1-infected and mock-infected mice. Mice with concomitant PMSS1 infection and active KRAS had greater expression of KI-67 and the SPEM marker CD44v10 (orthologous to human CD44v9, referred to herein as CD44v) and less expression of the IM marker TFF3 ^14^. Here we tested whether strains that had evolved during colonization of the human stomach, D1 and C2, could elicit similar phenotypes. Mice were infected with D1 or the stock or adapted C2 strains prior to induction of active KRAS and euthanized at six weeks (**Figure S2**). The median stomach loads were similar among the different *H. pylori* strains (**Figure 4A**) and similar to what we previously reported for PMSS1 ^14^. Interestingly, titers of the C2 adapted strain were significantly greater than D1 titers. We noted that loads were lower after six weeks of infection than after one week (compare to **Figures 1B, 2A**), consistent with the onset of adaptive immunity to infection.

**Figure 4.**
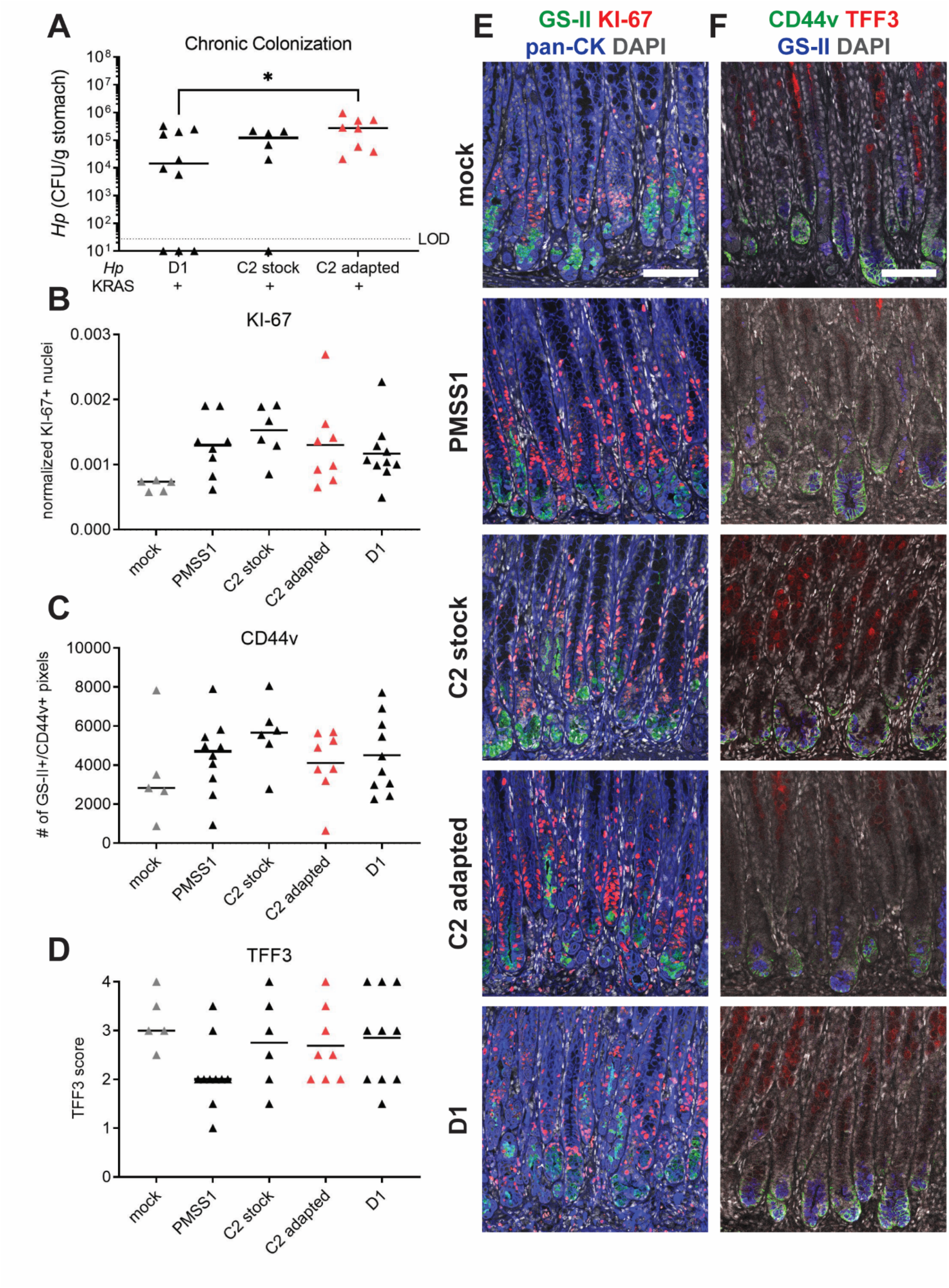
*H. pylori* isolates modulate preneoplastic progression in a strain-dependent manner. Mice were infected with the indicated *H. pylori* strains or mock-infected and active KRAS was induced. At six weeks, mice were euthanized and stomachs were assessed. (**A**) Stomach loads are shown. Data points represent actual values from each individual mouse, bars represent the median, and mice with no detectable CFU are plotted below the limit of detection (LOD). (**B-F**) IHC was used to detect expression of the indicated disease markers in thin stomach sections. Data are from N=2-3 independent experiments. (**B-D**) Three to five representative images per mouse were quantitatively or semi-quantitatively assessed and the median value for each mouse is plotted. Bars on the graphs indicate the median value for each treatment group. (**B**) KI-67+/DAPI+ nuclei were enumerated and normalized to the DAPI content (total number of DAPI+ pixels) of each image. (**C**) The number of GS-II+/CD44v+ pixels per image was quantified. (**D**) TFF3 staining was semi-quantitatively scored in a blinded fashion. (**E-F**) Representative images are shown. Scale bars, 100 µm. (**E**) Stomachs were stained with antibodies against KI-67 (red) and pan-cytokeratin (blue), the lectin GS-II (green) and DAPI (grey). (**F**) Stomachs were stained with antibodies against CD44v (green) and TFF3 (red), the lectin GS-II (blue) and DAPI (grey).

We next used immunohistochemistry (IHC) to assess the expression of KI-67, CD44v and TFF3 in mouse stomachs. For comparison, we also tested stomachs with PMSS1 or mock infection (from a previous study ^14^ and two additional replicates for this study). As we previously reported, KI-67 and CD44v expression were elevated in mice with PMSS1 infection compared to mock-infected mice, whereas TFF3 expression was decreased (**Figure 4B-F**). The other *H. pylori* strains also caused elevated KI-67 and CD44v expression, demonstrating that these phenotypes may be a common response to *H. pylori* infection in the context of metaplasia. However, only *H. pylori* strain PMSS1 reduced TFF3 expression. Thus, some features of gastric preneoplastic progression are shared among different *H. pylori* strains, whereas other features may be strain-dependent.

### Sequence comparison of D1 and C2 strains reveals mutations in outer membrane proteins

Isolates from the J99 collection exhibited differential colonization patterns in KRAS+ and KRAS-mice (**Figure 2A**), despite these strains having relatively few polymorphisms compared to isolates from different individuals; that is to say, D1 is more genetically similar to C2 than it is to PMSS1, for example. In order to investigate potential genetic determinants of colonization, we used our previously reported whole genome sequencing data ^27^ to examine the 349 polymorphic loci between isolates D1 and C2. For this study, we focused on the 175 polymorphisms that resulted in alterations to amino acid sequence, which includes 123 nonsynonymous single nucleotide polymorphisms (SNPs) and 52 indels detected across 68 genes, which included outer membrane proteins (OMPs) and predicted OMPs.

The *H. pylori* genome is enriched for repetitive regions, particularly in the genes encoding OMPs, which allow for high rates of recombination ^29, 30^. Therefore, we predicted that many of these alterations reflect inter- or intragenomic recombination events or mis-mapping of short reads to paralogs with high sequence identity. We leveraged ClonalFrameML to bioinformatically predict genes with mutations predicted to be introduced through importation of divergent DNA ^31^. As expected, more than half of these alterations (65%) were predicted to have been introduced via recombination events and were detected across only 14 genes. The remaining 46 SNPs and 16 indels differentiating C2 and D1, which are not predicted to arise from recombination, are listed in **Tables S1 & S2**.

To further address the limitations of mapping short read data to extended repeats in the reference, we generated long read assemblies of isolate D1 using Pacific Biosciences (PacBio) single-molecule real-time (SMRT) sequencing technology ^32^. Long read assemblies revealed that some of these polymorphisms predicted to be introduced through recombination were a result of short reads misaligning to the reference strain in repetitive regions. However, 10 of the 14 genes did have mutations or significant structural differences that were detected using both short and long read data (**Table S3**). Six out of ten encode outer membrane proteins: *sabA, sabB, homA, hopJ, hopK and hopQ*. Genes jhp0440, jhp1031 and *hopQ* had significantly altered protein lengths. Isolate D1 had an alternate start site rendering the protein 17 amino acids shorter than the C2 HopQ protein.

### The sabB locus undergoes dynamic gene conversion events in vivo

The adapted C2 isolate, which colonized the metaplastic stomach more robustly than the C2 stock strain (**Figure 2B**), was also sequenced using the PacBio SMRT sequencing platform to test if genetic changes occurred during acute colonization of the metaplastic stomach environment. Notably, the only region that differed between the two strains was the C-terminus of the outer membrane protein SabB. The consensus sequence of C2 adapted differed from C2 stock at five positions all within a short 50 nucleotide region. Only one of these polymorphisms results in an amino acid change: a mutation from threonine, an amino acid with a polar side chain, to alanine, an amino acid with a hydrophobic side chain, at position 553 (**Figure 5A**, red asterisk). Further analysis of the individual C2 stock Illumina reads revealed that the five polymorphisms detected in C2 adapted strain were present at a low frequency, indicating that there may be two alleles present in this population (**Figure 5A**). The predominant allele (variant 1) is estimated to be present at a frequency of ∼80% compared to 20% frequency of the minor allele (variant 2). Similar analysis of the PacBio sequencing reads demonstrated that in the C2 adapted strain, variant 2 increased in abundance to ∼40-50% (**Figure 5A**). Therefore, we hypothesized that variant 2 was selected for during gastric infection.

**Figure 5.**
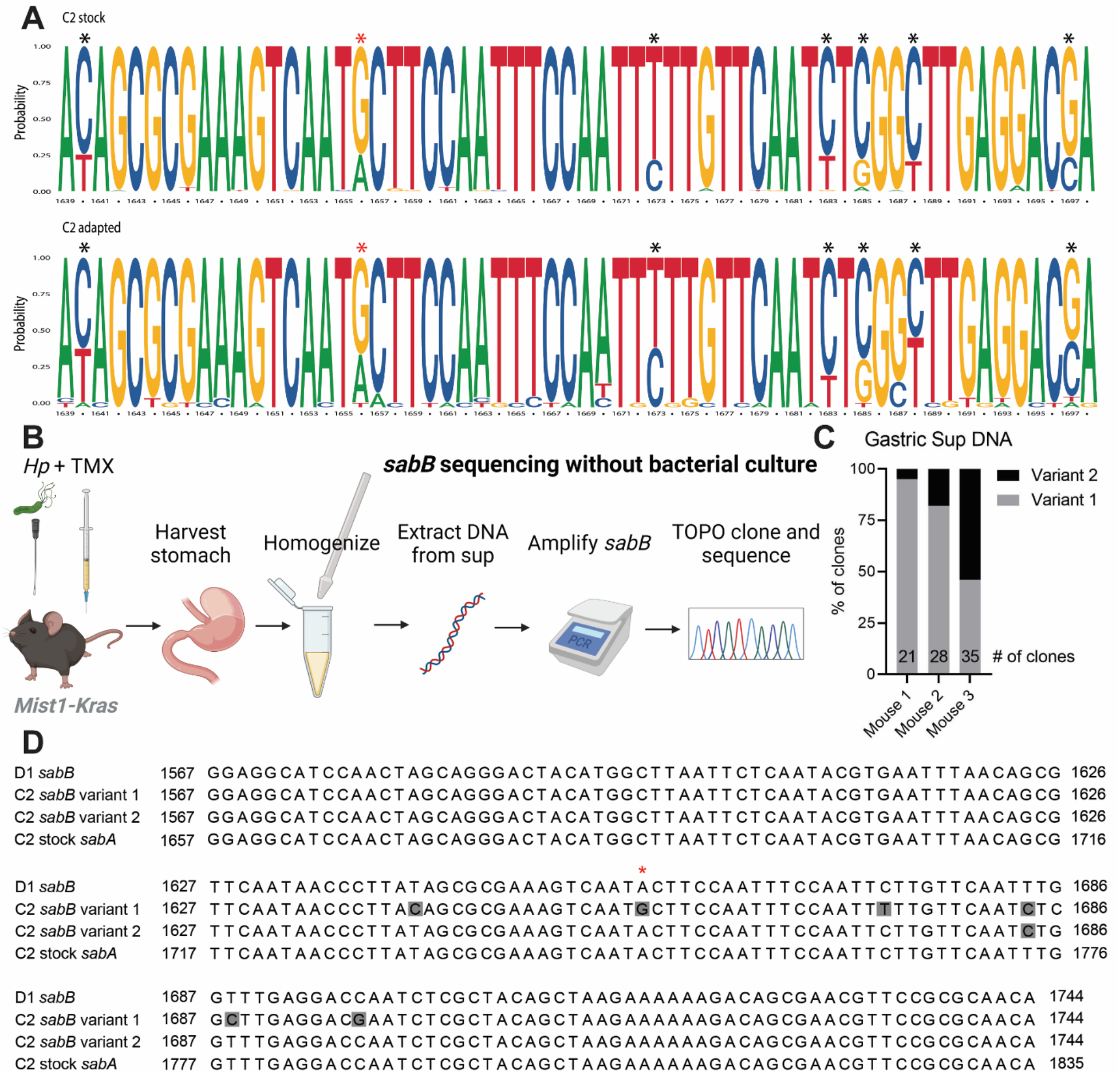
The C2 strain harbors two alleles of *sabB* that change in frequency during *in vivo* infection. (**A**) Sequence logos depict the probability of each base appearing at each position of the C-terminal region of *sabB* in C2 stock (top) and C2 adapted (bottom). Asterisks highlight positions with nucleotide variants. The red asterisk indicates the nonsynonymous change. (**B**) Schema for amplifying and sequencing *sabB* from mouse gastric homogenate supernatant. Illustration created with BioRender.com. (**C**) The proportion of TOPO clones of *sabB* variant 1 (grey) and variant 2 (black) is depicted for gastric homogenate supernatants from the indicated mice infected with C2 stock for six weeks. (**D**) The sequence alignment shows the C-terminal region of *sabB* in strains D1, C2 variant 1 and C2 variant 2, with nucleotide variants highlighted in grey. The red asterisk indicates a nonsynonymous change. The homologous region of *sabA* from C2 stock is also shown.

Attempts to use Sanger sequencing to detect the *sabB* variant 2 in genomic DNA isolated from either the C2 stock or adapted strains were unsuccessful, possibly because culturing the bacteria on plates prior to DNA extraction selected for variant 1. To circumvent this issue, we isolated DNA directly from gastric homogenate supernatants **(Figure 5B**) from mice infected with the C2 stock strain for six weeks (from **Figure 4A**). After PCR amplification of *sabB*, we used TOPO cloning and sequencing to test for the frequencies of variant 1 and 2 in each mouse. We found that the frequency of variant 2 ranged from 5 to ∼50% in different mice, confirming there are two alleles of *sabB* present in the C2 stock and that this genetic variation is dynamic during *in vivo* colonization (**Figure 5C**).

SabB has paralogs SabA and HopQ, which facilitate binding to sialic acid and CEACAMs, respectively ^33, 34^. Gene conversion events among these adhesins and putative adhesins are thought to modulate adherence to inflamed tissue during chronic infection ^35, 36^. All five SNPs in C2 *sabB* variant 2 were shared with the homologous region of *sabA*, suggesting that a recombination event gave rise to *sabB* variant 2 (**Figure 5D**). Strikingly, the *sabB* C-terminal region from the D1 strain also shared the five SNPs with C2 variant 2 and *sabA*, though D1 SabB was overall highly divergent from C2 SabB (**Figure S3**). Thus, variant 2 of *sabB* is associated with robust mouse infection and likely arose from a *sabA* recombination.

### sabB promotes tissue adherence and colonization of the mouse stomach

Because our *in silico* analysis suggested that *sabB* may be important during infection, we investigated this gene further. To test whether SabB, like its paralogues, could promote adherence of *H. pylori* to gastric tissue, we generated Δ*sabB* mutants in the C2 adapted and D1 strains and tested their ability to adhere to *ex vivo* gastric tissue compared to the parental strains. In KRAS-gastric tissue, adherence of both parental strains was greater than adherence of the isogenic Δ*sabB* mutants, both in the corpus and the antrum (**Figure 6A**). The same was true in KRAS+ stomachs (**Figure 6B**), and notably, the parental strains bound about twice as well to KRAS+ tissue than to KRAS-tissue. Interestingly, the C2 adapted strain was more adherent than D1 in KRAS-tissue, but the two strains were similar in KRAS+ tissue. We also generated a Δ*sabB* mutation in a derivative of G27, a highly tractable model *H. pylori* strain ^37^. Loss of *sabB* in this strain background also significantly reduced adherence to KRAS+ corpus and antrum tissue, and the defect could be rescued by expressing *sabB* at a neutral intergenic locus ^38^ (**Figure 6C**).

**Figure 6.**
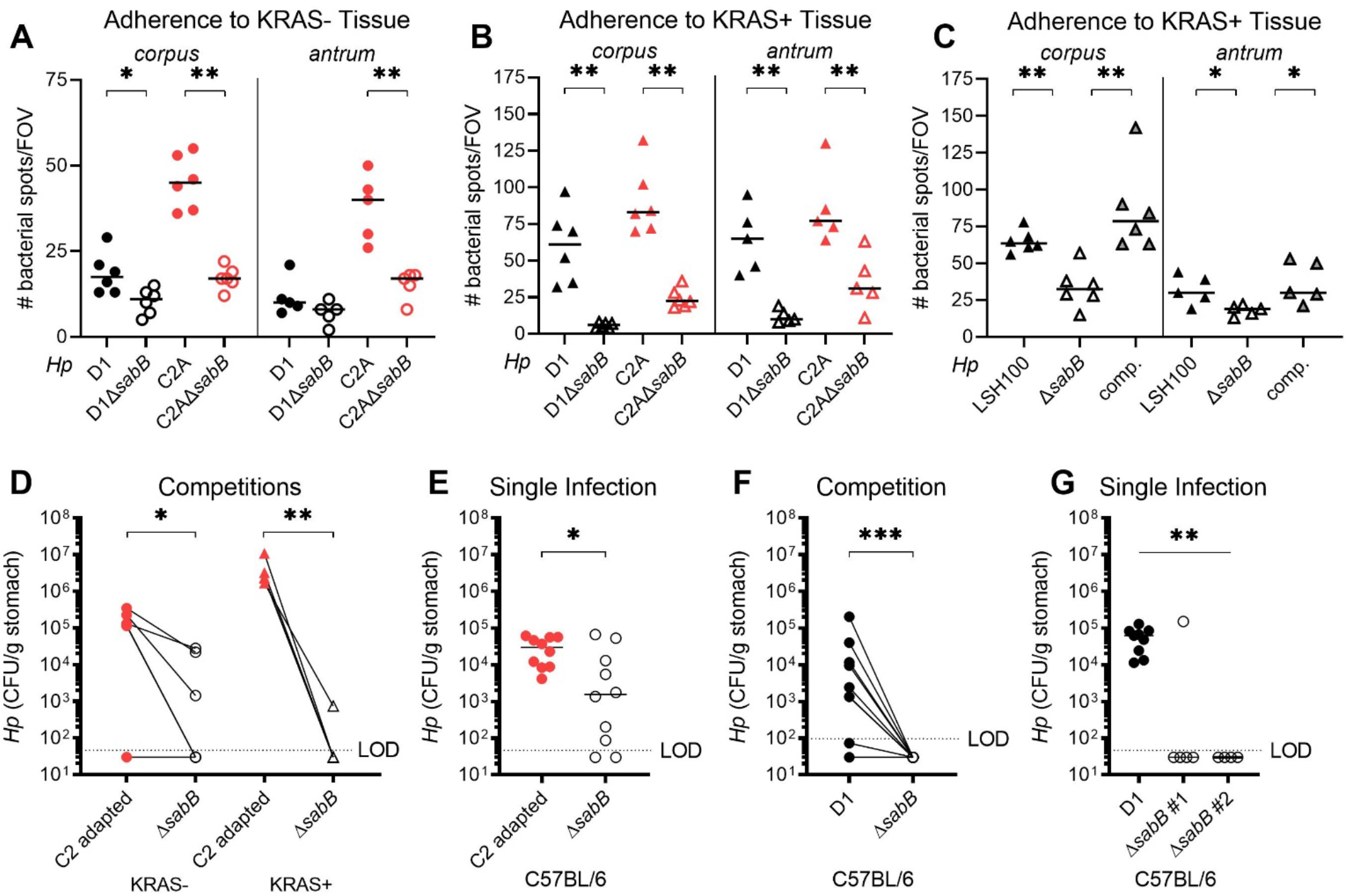
SabB promotes colonization of healthy and metaplastic stomach by the D1 and C2 adapted strains. (**A-C**) Tissue adherence was assessed using a previously described *ex vivo* binding assay ^36^. The indicated *H. pylori* strains were labeled with fluorescein isothiocyanate (FITC), applied to mouse tissue sections and the number of bacterial spots per field of view (FOV) was quantified. Each data point indicates one FOV and bars indicate the median value. Two to three technical replicates were performed with bacteria labeled on two different days and a representative experiment is shown. (**A**) The indicated strains were adhered to KRAS-tissue. C2A, the C2 adapted strain. (**B-C**) The indicated strains were adhered to KRAS+ tissue. (**C**) A *sabB* mutation (Δ*sabB*) was generated in a derivative of G27 (LSH100), and complemented (“comp.”) at a neutral intergenic locus. (**D-F**) Single infections and competitions were performed to compare wild-type and isogenic Δ*sabB* strains. (**C**) In *Mist1-Kras* mice, active KRAS was induced or sham-induced. Eight weeks later, mice were infected with a 1:1 mixture of the adapted C2 strain and Δ*sabB* mutant for one week. (**D-F**) C57BL/6 mice were infected with the indicated strains for one week. Data are from N=2 independent experiments. Data points represent actual values from each individual mouse, bars represent the median, and mice with no detectable CFU are plotted below the limit of detection. In **C** and **E**, lines connect wild-type and mutant titers from the same mouse. * *P* < 0.05, ** *P* < 0.01, *** *P* < 0.001. For **A-F**, statistical significance was assessed with a Mann-Whitney U test; for **F**, significance was assessed with a Kruskal-Wallis test.

Finally, to test whether SabB is necessary *in vivo*, we infected mice with the Δ*sabB* mutants and/or parental strains. First, eight weeks after active KRAS induction or sham induction, mice were infected with a 1:1 mixture of the C2 adapted strain and Δ*sabB* mutant (**Figure 6D**). The Δ*sabB* mutant was somewhat attenuated in KRAS-mice and highly attenuated in KRAS+ mice, demonstrating that SabB promotes gastric colonization. Because the Δ*sabB* mutant had a phenotype in KRAS- (i.e. healthy) mice, we used C57BL/6 mice to further probe the role of SabB in gastric infection. The C2 adapted Δ*sabB* mutant performed worse than the parental strain in single infection (**Figure 6E**), demonstrating that its *in vivo* attenuation occurs whether or not the wild-type strain is present. Strikingly, the D1Δ*sabB* strain was not recovered from any mice in competitive infections (**Figure 6F**) and only from one out of five mice in single infections (**Figure 6G**). A second clone was not recovered from any mice (**Figure 6G**), which suggests that the colonization defect was due to the lack of *sabB*, not due to an off-target effect with the constructed strain. Taken together, these findings demonstrate that SabB promotes *H. pylori* adherence to tissue and is required for infection.

## Discussion

It has been well established that prolonged *H. pylori* infection can induce premalignant tissue changes, but little is known about the dynamics between host and pathogen at later stages of disease development. In this study we used the *Mist1-Kras* mouse model to examine *H. pylori* strain-specific determinants of stomach and gland colonization during the development of gastric metaplasia. We found that *H. pylori* strains have different propensities for infecting the stomach with and without metaplasia (+/-KRAS). Strains that can robustly colonize KRAS+ mice have an expanded niche in the corpus glands, similar to what is observed with long term (>4wks) chronic infections in mice ^8, 10^. The ability of *H. pylori* to infect KRAS+ mice can be selected for during acute infection of metaplastic tissue via changing allele frequencies of the putative adhesin *sabB*.

J99 strains, originating from the same patient at two different stages of disease ^25, 26^, exhibited differential colonization in KRAS+ mice. J99 is a relatively poor colonizer of both KRAS- and KRAS+ mice. Strain SC4, isolated at the early timepoint but belonging to a different genetic subgroup from J99, was also a relatively poor colonizer of KRAS +/-mice. Isolate D1, collected from the patient during a period of gastric atrophy, robustly colonized both KRAS- and KRAS+ mice, and titers were greater in KRAS+ stomachs than KRAS-stomachs. In contrast, isolate C2, originating from the same time point as D1, had relatively poor colonization in some mice but very robust colonization in others. We showed that C2 could be adapted to *in vivo* colonization through just one week of infection in the KRAS+ stomach by selecting for a lower frequency allele at the *sabB* locus.

Not all of the clinical isolates we tested colonized the metaplastic stomach of KRAS+ mice, indicating there are bacterial determinants of metaplastic colonization. The host environment where clinical strains were isolated from may make certain isolates better adapted for colonization. Complete genome sequencing suggests that changes in the outer membrane protein SabB facilitated quick adaptation to the metaplastic stomach environment in isolate C2. SabB is a protein that is part of a paralogous family including HopQ, which binds to CEACAMs, and SabA, which binds to sialic acid ^33, 34^. SabA was previously reported to mediate *H. pylori* interactions with SPEM (metaplastic) cells ^39^, suggesting that this protein family is an important tool in *H. pylori*’s toolbox for late disease-state persistence. These proteins are some of the most highly variable in the *H. pylori* genome and can recombine to quickly adapt to changes to the gastric environment ^36, 40^. The altered bases within *sabB* variant 2 align to the paralog *sabA* from the C2 stock strain, indicating that these changes were most likely introduced through an intra-genomic recombination event between *sabA* and *sabB*. Inflamed tissue is more likely to express sialylated glycoconjugates than healthy tissue is ^34^, which may suggest that *H. pylori* genetically alters *sabA* and *sabB* in order to modulate binding to inflamed tissue. Our data suggest that *sabB* may play a previously unknown role in achieving colonization of tissue that is inflamed or exhibiting SPEM. However, it is likely not the only player. Other differences between C2 and D1 include mutations in two fucosyltransferases, *fucA* and *fucU,* which are involved in modifying bacterial lipopolysaccharide to mimic surface sugars on the host epithelium in order to evade immune detection ^41^; and several mutations in genes involved in DNA uptake, metabolism, and repair including *comB8*, *comM*, *hsdR*, *hsdM*, and *hsdS*.

In humans, *H. pylori* triggers chronic gastritis. In a subset of individuals, this gastric inflammation leads to gastric atrophy (loss of acid-producing parietal cells), which can then progress to metaplasia, dysplasia, and finally cancer. Once this pathogenic cascade is initiated, it is thought that *H. pylori* does not further contribute to the development of cancer and that these tissue alterations are not favorable to *H. pylori* proliferation. This is referred to as the ‘hit and run’ theory of gastric cancer development ^42^. This implies eradication of *H. pylori* infection is unnecessary once metaplastic lesions have developed. However, a long-term study with a Colombian cohort shows that eradication of *H. pylori* after discovery of metaplastic legions reduces risk of gastric cancer, suggesting a new role for *H. pylori* in the development or acceleration of gastric cancer ^43^. Host-pathogen interactions during pre-malignant tissue alterations are understudied, but we recently showed that *H. pylori* interacts with tissue in ways that impact disease progression and skew the immune response ^14^. In another mouse model of gastric atrophy and SPEM, induced via chemical ablation of parietal cells, *H. pylori* was shown to preferentially interact with SPEM cells within the corpus glands ^39^. Notably, KRAS+ mice have moderate gastric inflammation marked by M2 macrophage expansion ^14, 19^. Here we show that certain strains of *H. pylori* readily colonize the metaplastic stomach despite this preexisting inflammatory milieu, which might have been predicted to reduce or even clear infection. Furthermore, some *H. pylori* strains exhibited an expanded niche in the corpus glands. Together these data suggest that metaplastic environments may be suitable or even preferable for *H. pylori* growth and challenge the hit-and-run theory of carcinogenesis. This new model provides avenues to further explore *H. pylori* interactions with metaplastic tissue and cell types specific to metaplastic glands.

It is remains unknown if *H. pylori* has evolved to exploit pre-neoplastic tissue changes such as SPEM, IM, and dysplasia in order to expand its niche and increase transmission, or if pre-neoplasia is an unintended consequence of bacterial growth that is tolerated by, but not advantageous to, *H. pylori*. However, our findings suggest that certain bacterial strain-specific factors may promote colonization of the metaplastic stomach and these factors can vary among isolates in a single infected individual during the course of disease development. Additional studies of bacterial genetic factors facilitating colonization during SPEM may identify important predictors for risk of transmission and/or disease progression.

## Materials and Methods

### Ethics Statement

All mouse experiments were performed in accordance with the recommendations in the National Institutes of Health Guide for the Care and Use of Laboratory Animals. The Fred Hutchinson Cancer Center is fully accredited by the Association for Assessment and Accreditation of Laboratory Animal Care and complies with the United States Department of Agriculture, Public Health Service, Washington State, and local area animal welfare regulations. Experiments were approved by the Fred Hutch Institutional Animal Care and Use Committee, protocol number 1531.

### H. pylori *culture*

The *H. pylori* strains used in this study were PMSS1 ^20^, 7.13 ^44^, the G27 ^45^ derivatives NSH57 ^23^ and LSH100 ^37^, and four representative isolates from the J99 culture collection ^25, 27, 29^: J99, SC4, D1 and C2. All *H. pylori* isolates were grown on solid media, horse blood agar (HB agar). HB agar plates contain 4% Columbia agar base (Oxoid), 5% defibrinated horse blood (Hemostat Labs), 10 mg/ml vancomycin (Thermo Fisher), 2.5 U/ml polymyxin B (Sigma-Aldrich), 8 mg/ml amphotericin B (Sigma-Aldrich), and 0.2% β-cylodextrin (Thermo Fisher). For HB agar plates used to grow *H. pylori* from homogenized mouse stomach, 5 mg/l cefsulodin (Thermo Fisher), 5 mg/l trimethoprim (Sigma) and 0.2 mg/ml of bacitracin (Acros Organics, Fisher) were added to prevent outgrowth of mouse microflora. For competition experiments, 15 µg/ml chloramphenicol was added to distinguish between mutant (chloramphenicol-resistant) and wild-type (chloramphenicol-sensitive) bacteria. Shaking liquid cultures were grown in Brucella broth (Thermo Fisher) supplemented with 10% heat-inactivated FBS (Gemini BioProducts). *H. pylori* on plates and in liquid culture were grown at 37°C in a micro-aerobic conditions (10% CO_2_, 10% O_2_, and 80% N_2_) using a tri-gas incubator.

### Mist1-Kras *mouse model*

All experiments used Mist1-CreERT2 Tg/+; LSL-K-RAS G12D Tg/+ (“*Mist1-Kras*”) mice described previously ^19^ or C57BL/6J mice as indicated. Mice were housed in sterilized microisolator cages with irradiated rodent chow, autoclaved corn cob bedding, and acidified, reverse-osmosis purified water. Mice were genotyped from ear punches as previously described^19^. Expression of the KRAS transgene was induced via subcutaneous administration 5 mg of tamoxifen (Sigma Aldrich) in corn oil (Sigma Aldrich) for three consecutive days. Sham induced mice were administered corn oil subcutaneously on the same schedule. All mouse experiments were performed as previously described ^14^. The inoculum for each infection was 5 x 10^7^ cells of the indicated strain or strains. After stomach excision, the forestomach was removed and the stomach was opened along the lesser curvature. Stomachs were divided into equal thirds or halves containing both antral and corpus regions. For culture, stomach portions were placed in 0.5 mL of sterile BB10 media, weighed, and homogenized. Serial homogenate dilutions were plated on nonselective HB plates, or both nonselective and chloramphenicol-containing plates for competition experiments. Homogenates were then pelleted at 15,000 rpm in a benchtop centrifuge (Eppendorf 5424) and the supernatant was stored at −20^°^C. After 5-9 days in tri-gas incubator, colony-forming units (CFU) were enumerated and reported as CFU per gram of stomach tissue.

### Gland occupation analysis

PMSS1 gland colonization was assessed as previously described ^8^ with the following modifications. One third of the stomach was embedded in 4% agarose in 1x phosphate-buffered saline (PBS, Gibco) and cut into 200 μm thick longitudinal sections using a Leica VT1200S Vibratome. Tissue sections were then permeabilized overnight at 4^°^C by gently rocking in blocking buffer comprising 3% bovine serum albumin (Sigma Aldrich), 1% saponin (Sigma Aldrich) and 1% Triton X-100 (Sigma Aldrich) in phosphate buffered saline (PBS). Stomachs were incubated with 1:1,000 rabbit polyclonal anti-*H. pylori* PMSS1 antibody (gift of Manuel Amieva, Stanford University) and 1:2,000 GS-II 488 (conjugated lectin from *Griffonia simplicifolia*, Fisher) for two hours at 4^°^C with gentle rocking. After three ten-minute washes in PBS, samples were incubated with 1:2,000 Alexa Fluor 647 donkey anti-rabbit IgG (Invitrogen) and 1:2,000 DAPI for two hours at room temperature with gentle rocking. After three ten-minute washes in PBS, sections were mounted onto glass slides with imaging spacers cut from Parafilm (Bemis) in ProLong Diamond Anti-Fade Reagent (Molecular Probes) and coverslips were sealed with VaLP (1:1:1 Vaseline:Lanolin:Paraffin). Stomach sections were imaged on a Zeiss LSM 780 laser-scanning confocal microscope, or with an UltraView spinning disk microscope (PerkinElmer). Z-stacks were collected to visualize *H. pylori* within glands and assessed in Volocity (Quorum Technologies) to enumerate *H. pylori* in glands based on fluorescent voxels as previously described ^8^.

### Immunohistochemistry to assess tissue markers

Immunohistochemistry was performed as previously described ^14^. Briefly, thin sections of formalin-fixed, paraffin-embedded tissue were deparaffinized with Histo-Clear solution (National Diagnostics) and rehydrated in decreasing concentrations of ethanol. Slides were boiled in a pressure cooker for 15 min in Target Retrieval Solution (Agilent Dako) for antigen retrieval. Slides were incubated with Protein Block, Serum Free (Agilent Dako) for 90 min at room temperature. Primary antibodies (**Supplementary Table S4**) were diluted in Protein Block, Serum Free, or Antibody Diluent, Background Reducing (Agilent Dako), and applied to the slides overnight at 4°C. Secondary antibodies were diluted 1:500 in Protein Block, Serum Free and slides were incubated for one hour at room temperature protected from light. Slides were mounted in ProLong Diamond antifade reagent with DAPI (Invitrogen) and allowed to cure for 24 hours at room temperature before imaging. Slides were imaged on a Zeiss LSM 780 laser-scanning confocal microscope using Zen software (Zeiss). For assessment of *H. pylori* gland colonization, the entire length of each corpus was inspected for *H. pylori* cells and three to five representative images per sample were taken. For assessment of epithelial disease markers, three to five representative images of the corpus were taken per sample.

### PacBio long read sequencing

Single molecule real-time sequencing (SMRT-Seq) was carried out on a PacBio Sequel-I instrument (Pacific Biosciences, USA). Genomic DNA to be sequenced was purified using the Wizard Genomic DNA Purification Kit (Promega), concentration was determined using the Qubit dsDNA HS (High Sensitivity) assay kit (Thermo Fisher), and purity was calculated using a NanoDrop One (Thermo Fisher). Genomic DNA samples (3 µg) were sheared to an average size of 12 kb via G-tube (Covaris) before library preparation. Libraries were then generated with SMRTbell Express Template Prep Kit 2.0 and pooled libraries were size-selected using the BluePippin system (Sage Sciences) at a 4 kb minimum threshold. Sequencing reads for the D1 strain were processed using the Pacific Biosciences SMRTAnalysis pipeline version 8.0.0.80529 and assembled using Microbial Assembler. Genome assembly showed 21,312 polymerase reads that were further partitioned into 210,582 subreads with an N50 value of 5,441 nucleotides and a total number of subread bases of 767,735,588 with a mean coverage of 397x. Genome assembly of D1 resulted in a single contig: a chromosomal sequence of 1,685,094 bp. Sequencing reads for the C2 adapted strain were processed using the Pacific Biosciences SMRTAnalysis pipeline version 8.0.0.80529 and assembled using Flye de novo assembler ^46^. Genome assembly showed 5,657 polymerase reads that were further partitioned into 41,337 subreads with an N50 value of 6,850 nucleotides and a total number of subread bases of 141,837,408 with a mean coverage of 69x. Genome assembly of the C2 adapted strain resulted in a single contig: a chromosomal sequence of 1,653,204 bp.

### Bioinformatics

Short reads for isolates C2 stock and D1 were downloaded from the NCBI SRA database (BioProject accession: PRJNA622860). Single nucleotide variants differentiating D1 and C2 stock were determined by aligning short reads to J99 reference (AE001439) with BreSeq v0.35.0 software using default parameters ^47^. ClonalFrameML was used to predict putative sites of recombination ^31^. The D1 and C2 adapted isolates were sequenced on a PacBio Sequel instrument to generate long sequence reads. Closed reference sequences were generated using either SMRT Link web-based analysis suite <https://www.pacb.com/support/software-downloads/> or Flye <https://github.com/fenderglass/Flye> assembly pipelines and genomes were annotated using Pathosystems Resource Integration Center (PATRIC, <https://www.patricbrc.org>) annotation tool with *Helicobacter pylori* (species ID: 210) as the reference. Recombination events predicted by short read assemblies were validated by cross comparing with sequences generated from long read assemblies. Genomic comparisons of C2 stock vs. adapted isolates were performed by aligning C2 stock short reads with Breseq v0.35.0. using C2 adapted PacBio assembly as the reference. The frequency matrix of each base at each position for the short read sequencing was generated using pileup2acgt from the Sequenza package on the previously reported SAMtools pileup ^27, 48, 49^. The frequency matrix for the long read data was generated by mapping the reads to the Flye assembly using Minimap2 ^50^ and running the output through bam-readcount <https://github.com/genome/bam-readcount>. The sequence logos were generated with ggseqlogo <htttps://CRAN.R-project.org/package==ggseqlogo> from ggplot2 ^51^.

### TOPO cloning and sequencing

DNA was isolated from stored mouse gastric homogenate supernatants via enzymatic digestion and phenol/chloroform extraction ^52^. The C-terminal region of *sabB* was amplified by PCR using primers 5’aagctcaaggcaatctctgtgc3’ (sabBFor) and 5’gatcatgcgtttttgatccctgg3’ (jhp0660R) ^36^. The PCR product was verified by agarose gel electrophoresis and used for TOPO cloning with the Zero Blunt® TOPO® PCR Cloning Kit (Invitrogen). After selection on kanamycin, clones were screened by colony PCR with sabBFor and jhp0660R primers and Go Taq master mix (Promega). Dye-terminator sequencing using BigDye (Thermo Fisher) with the jhp0660R primer was performed by the Fred Hutch Genomics Shared Resource and the results were analyzed using SnapGene software version 5.2.4 (Insightful Science).

### Construction of H. pylori mutants

Deletion mutants of *sabB* (Δ*sabB*) were constructed in four *H. pylori* strain backgrounds (D1, C2 adapted, LSH100 *rdxA::aphA3sacB*, and LSH100 *hp0203-0204* intergenic region::*sabB*). Deletions of *sabB* from D1 and C2 adapted were generated by transforming parent strains with gDNA from J99 *sabB::catsacB* ^36^. The gDNA was extracted using a Wizard Genomic DNA Purification Kit (Promega). Clones were selected on HB plates with 15 µg/ml chloramphenicol as previously described ^53, 54^. Clones were confirmed using diagnostic PCR and Sanger sequencing with primers 5’ tgggttgagatcatgcaagcat 3’ (jhp0658F) and 5’ gatcatgcgtttttgatccctgg 3’ (jhp0660R) ^36^. LSH100 *rdxA::aphA3sacB sabB::catsacB* was generated with the same strategy but using gDNA extracted from D1Δ*sabB*. Strains were back-crossed to reduce the likelihood of off-target mutations. To complement the mutation, *sabB* was amplified from strain D1 using primers McGee_sabB_fwd (5’ tagaactagtggatccattttcatttctattcatgtttacaataaaaaaattactttaag 3’) and McGee_sabB_rev (5’ atcgataagcgaattcttaataagcaaacacataattgagatacacgctataaagc 3’) and cloned into the pDYC40 plasmid that contains a kanamycin resistance cassette ^55^ via In-Fusion cloning (Takara). This plasmid is designed for complementation at a previously characterized, neutral intergenic chromosomal site (*hp0203-0204* intergenic region) ^38^. LSH100 was transformed with the resulting pDYC40::*sabB* plasmid and clones were selected on HB agar with 25 µg/ml kanamycin. The native *sabB* locus was then mutated by transforming with a PCR product prepared from D1Δ*sabB* using primers jhp0658F and jhp0660R to amplify the *catsacB*- interrupted *sabB* region and selecting on HB agar with 15 µg/ml chloramphenicol. All mutants were confirmed with PCR and Sanger sequencing.

### Ex vivo tissue adherence assay

The *ex vivo* tissue adherence assay was performed as previously described ^36^. Briefly, bacteria used for the assay were grown in 10-20 mL of BB10 (10% FBS in Brucella broth) overnight to achieve an optical density at 600 nm (OD_600_) between 0.5 and 1.0. Bacteria were pelleted by centrifuging and then washed in PBS-T (0.05% Tween 20 in phosphate-buffered saline). Bacteria were resuspended in carbonate buffer and incubated with fluorescein 5(6)-isothiocyanate (FITC, Sigma) dissolved in DMSO (dimethyl sulfoxide, Alfa Aesar) for 10 minutes and washed in oxidized 1% bovine serum albumin (BSA). The labeled organisms were stored in oxidized 1% BSA at −80°C. Slides with mouse stomach tissue sections were deparaffinized in Histo-Clear solution (National Diagnostics) and rehydrated with isopropanol, ethanol, deionized water, and PBS. Slides were blocked with oxidized 1% BSA for two and a half hours in a hydration chamber before incubating with FITC-labeled bacteria (OD_600_ = 0.01) for another two hours. The excess bacteria were washed off in PBS-T and the slides were mounted in ProLong Diamond antifade reagent with DAPI (Invitrogen). The slides were allowed to cure for 24 hours at room temperature before imaging at 400x magnification on a Zeiss LSM 780 laser-scanning confocal microscope using Zen software (Zeiss). For the quantification of FITC-labeled bacteria adhered to the tissue, four to six representative images per antrum or corpus were analyzed using FIJI software (National Institutes of Health) to count the number of bacterial spots in each image field.

### Statistical analysis

Statistical analyses were performed according to the tests specified in each figure legend using Prism v9 software (GraphPad). P-values less than or equal to 0.05 were considered statistically significant and are marked with asterisks ( *, *P* < 0.05; **, *P* < 0.01; *** *P* < 0.001).

### Data availability

The previously published Illumina sequencing reads for the C2 stock strain ^27^ are available at the NCBI SRA database database (BioProject accession number PRJNA633860), <https://www.ncbi.nlm.nih.gov/sra/PRJNA622860>. The complete genomes assembled from the PacBio SMRT sequencing performed in this study are available at NCBI GenBank under Bioproject PRJNA786001.

## Acknowledgements

The authors wish to thank Dr. Richard M. Peek, Jr. at Vanderbilt University for sharing the *H. pylori* J99 strain collection. This work was funded by an Innovation Grant from the Pathogen-Associated Malignancies Integrated Research Center at Fred Hutchinson Cancer Research Center and NIH R01 AI54423 to NRS and NIH R01 DE027850 to CDJ. Research was supported by the Cellular Imaging, Comparative Medicine, and Genomics & Bioinformatics of the Fred Hutch/University of Washington Cancer Consortium (P30 CA015704). VPO is a Cancer Research Institute Irvington Fellow supported by the Cancer Research Institute, and was also supported by a Debbie’s Dream Foundation—AACR Gastric Cancer Research Fellowship, in memory of Sally Mandel (18-40-41-OBRI). LKJ was supported by the Chromosome Metabolism and Cancer Training Grant (NCI T32 CA009657). Some studies used a microscope that was purchased with a grant from the M. J. Murdock Charitable Trust.

## Supporting Materials

**Supplemental Movie 1.** Shown is a Z-stack of a mouse stomach infected with *H. pylori* strain PMSS1(green) for one week, 12 weeks after tamoxifen was given to induce active KRAS. DAPI, blue, depicts nuclei. GS-II, red, labels metaplastic cells at the base of the glands. The tissue was imaged at 100x magnification on a Zeiss LSM 780 confocal microscope.

**Figure S1.**
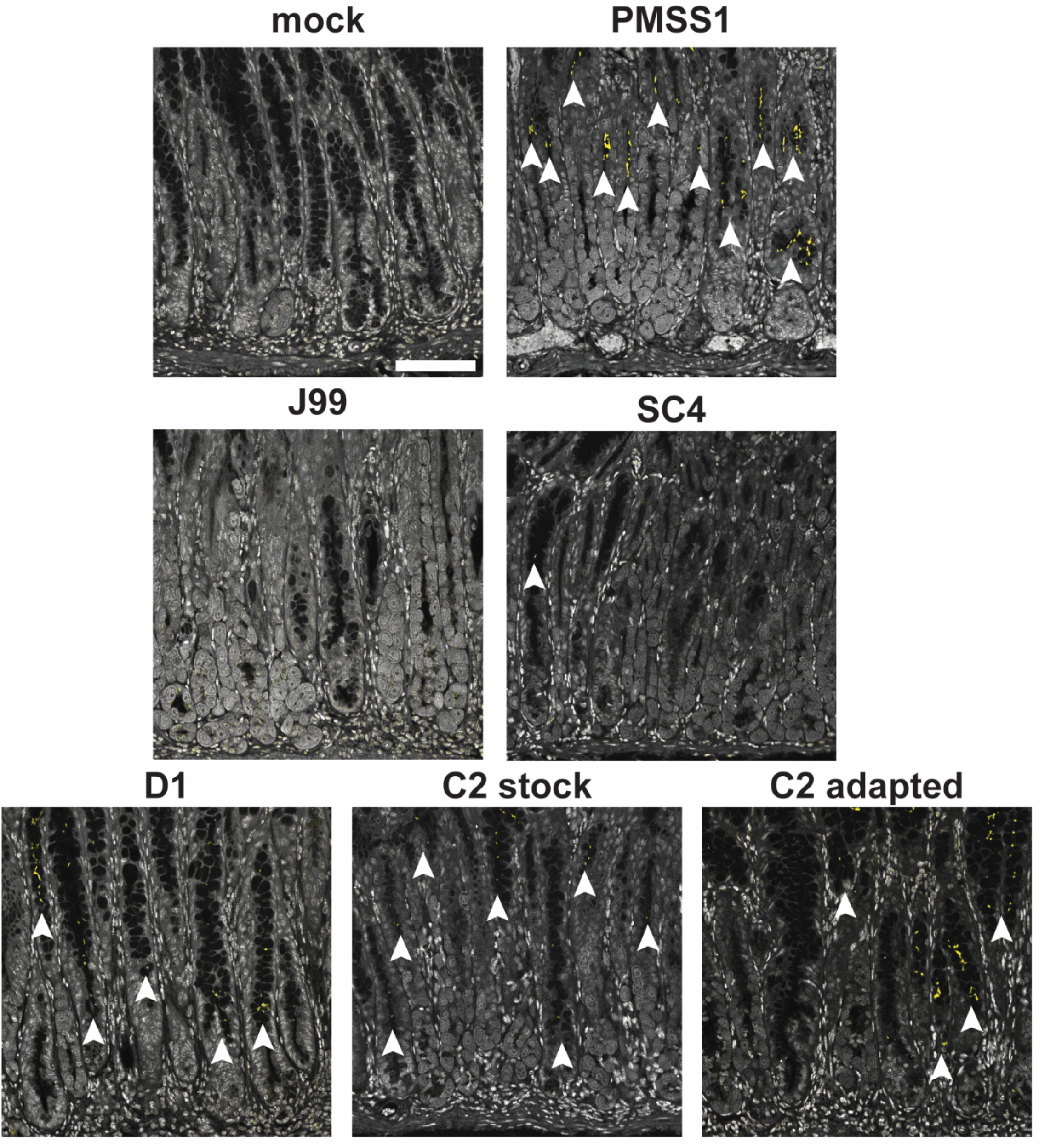
Different *H. pylori* isolates have different propensities for colonizing KRAS+ glands. Representative images of corpus tissue are shown. Grey indicates nuclei and yellow indicates *H. pylori* cells. Arrowheads show examples of bacteria within glands. Scale bar, 100 µm.

**Figure S2.**
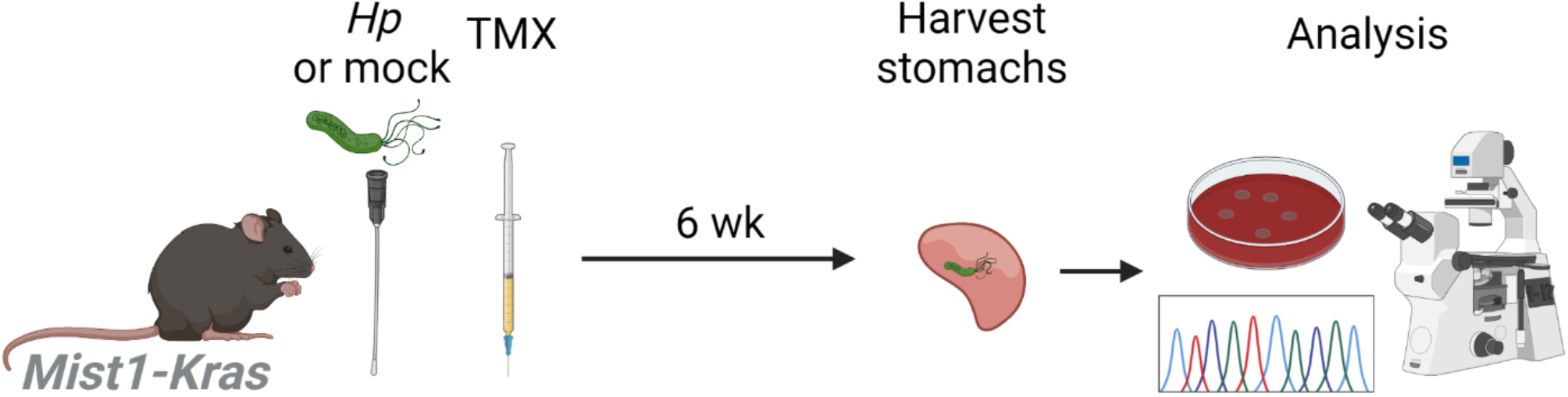
Mouse model for assessing whether *H. pylori* strains impact metaplasia development. Mice are initially infected with various *H. pylori* strains or mock-infected. All mice receive tamoxifen (TMX) to induce active KRAS expression. After six weeks, mice are humanely euthanized. Stomachs are harvested and assessed by immunofluorescence microscopy analysis of disease marker expression, as well as *H. pylori* culturing and sequencing. Illustration created with BioRender.com.

**Figure S3.**
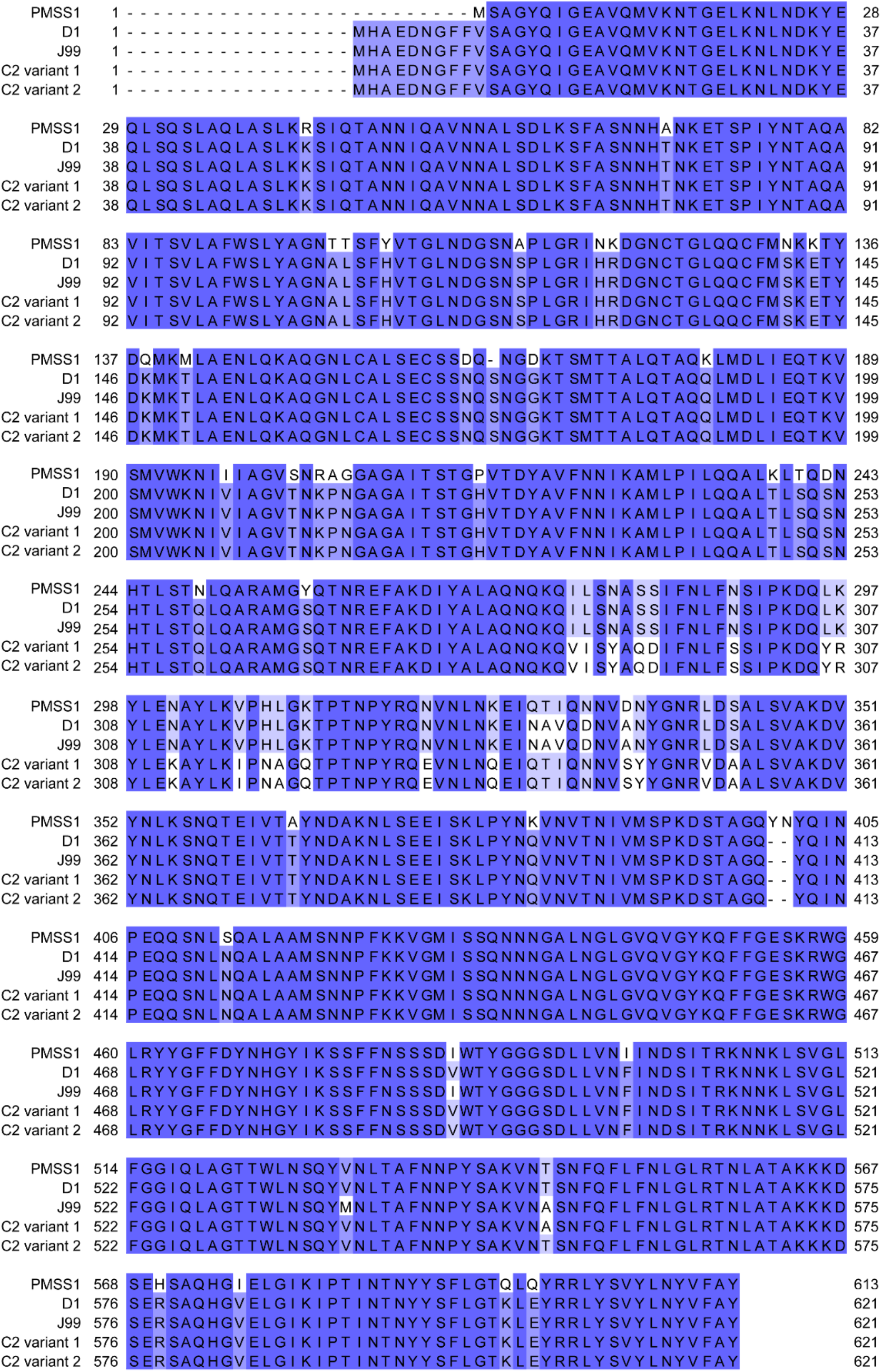
The outer membrane protein SabB differs between PMSS1 and strains from the J99 collection. Shown is a sequence alignment of the SabB protein comparing PMSS1 with D1, J99, and C2 variants 1 and 2. Amino acid changes are indicated.

**Table S1.**
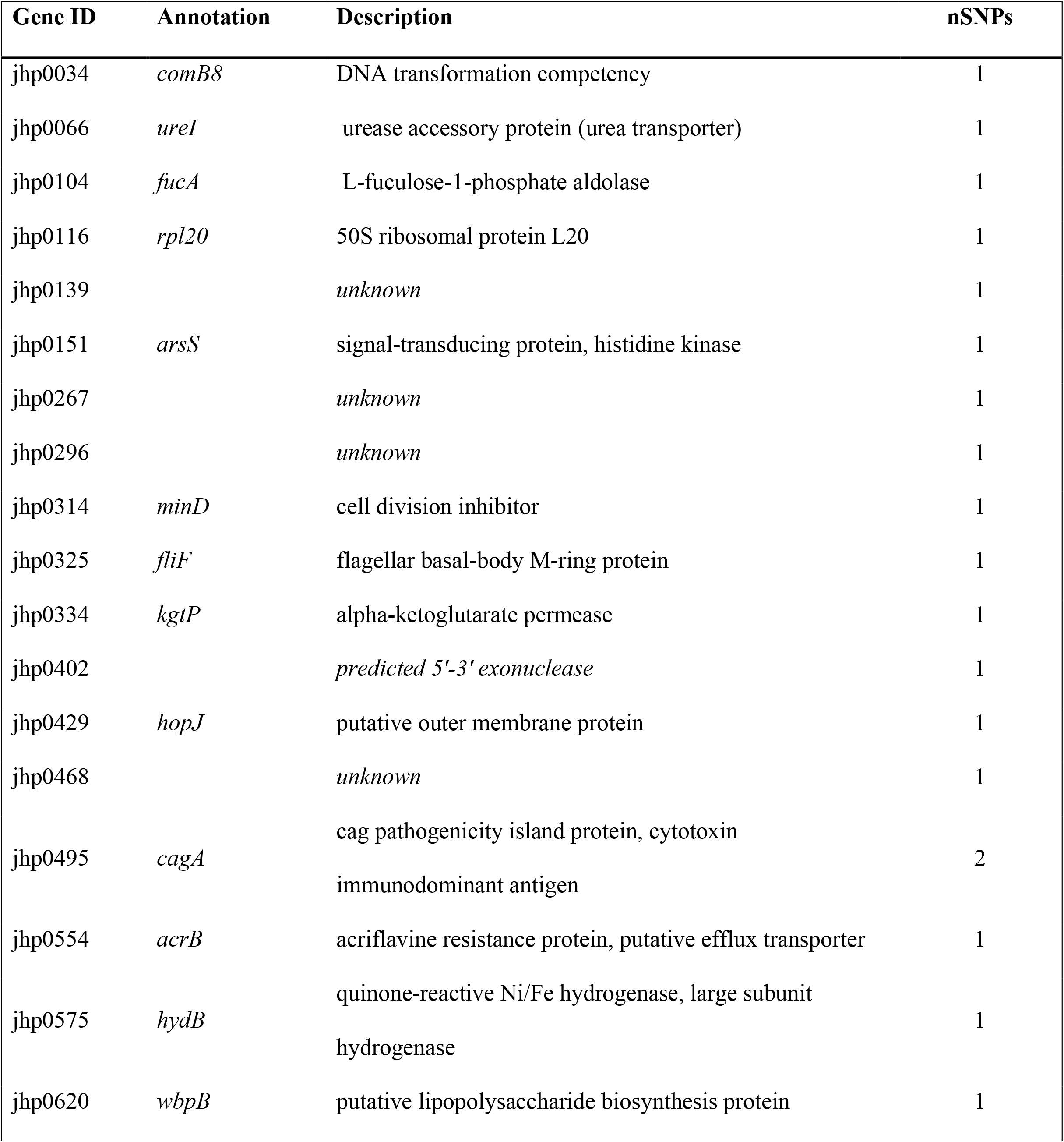
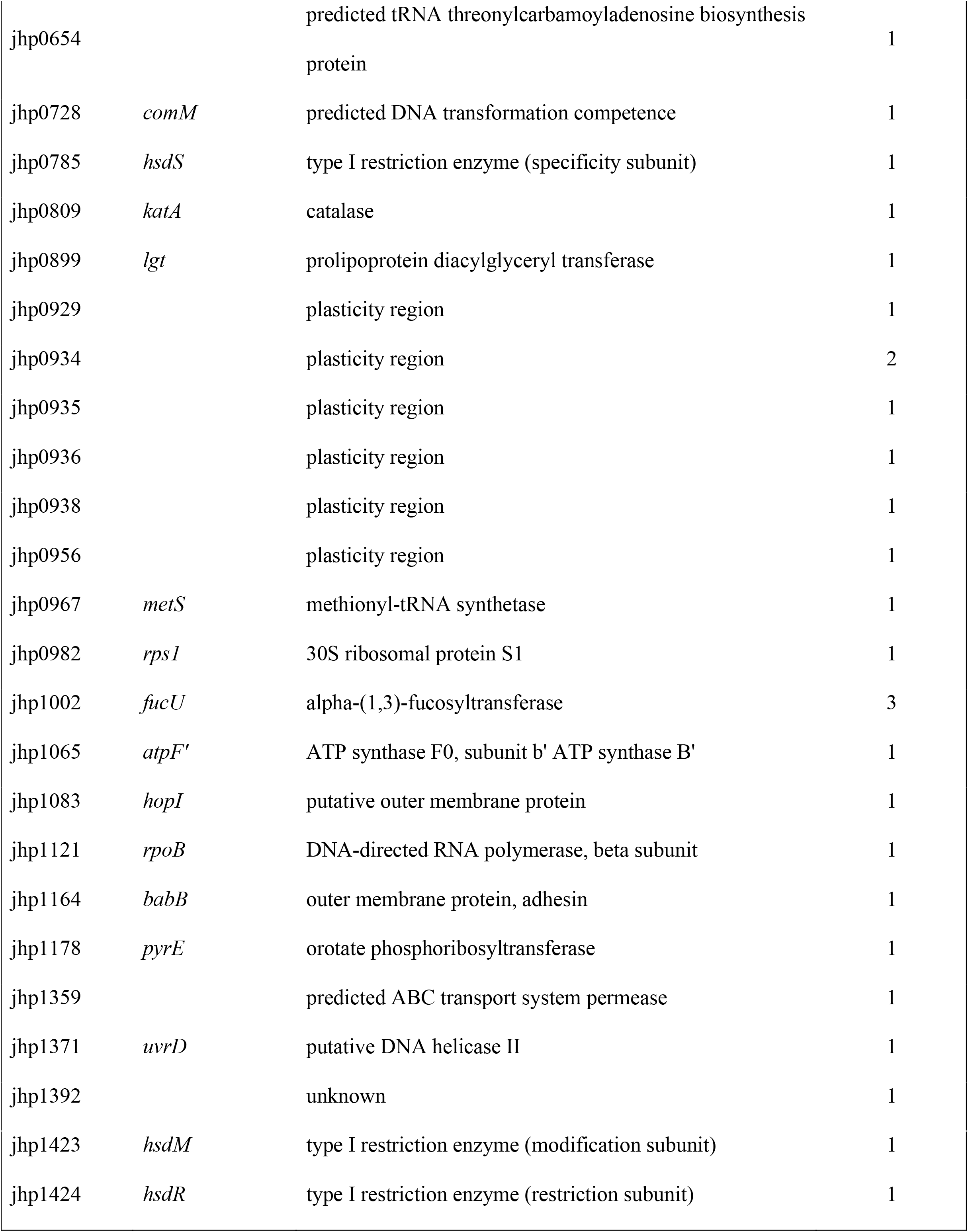
Nonsynonymous SNPs (n=46) between isolates D1 and C2 stock.

**Table S2.** Insertion or deletion events (n=16) between isolates D1 and C2 stock.

| **Gene ID** | **Annotation** | **Description** | **Indels** |
| --- | --- | --- | --- |
| jhp0151 | *arsS* | signal-transducing protein, histidine kinase | 1 |
| jhp0241 |  | conserved hypothetical secreted protein | 1 |
| jhp0451 | *pldA* | putative phospholipase A1 precursor (DR-phospholipase A) | 1 |
| jhp0709 | *amiA* | putative N-acetylmuramoyl-L-alanine amidase | 1 |
| jhp0741 | *lex2B* | Lipoligosaccharide 5G8 epitope biosynthesis associated protein | 1 |
| jhp0888 |  | oxygen-insensitive NAD(P)H nitroreductase | 1 |
| jhp0892 |  | unknown | 1 |
| jhp0928 |  | plasticity region | 1 |
| jhp0934 |  | plasticity region | 1 |
| jhp0937 |  | plasticity region | 1 |
| jhp0938 |  | plasticity region | 1 |
| jhp1002 | *fucU* | alpha-(1,3)-fucosyltransferase | 1 |
| jhp1171 | *csd5* | Cell-shape determining protein | 1 |
| jhp1305 |  | unknown | 1 |
| jhp1312 |  | unknown | 1 |
| jhp1410 |  | type III restriction enzyme (restriction enzyme) | 1 |

**Table S3.** Variation between C2 and D1 predicted to be the result of recombination.

| **Gene ID** | **Annotation** | **SNPs** | **nSNPs** | **Indels** |
| --- | --- | --- | --- | --- |
| jhp1300 |  | 2 | 2 | 3 |
| jhp0659 | *sabB* | 13 | 6 | 0 |
| jhp0662 | *sabA* | 2 | 1 | 1 |
| jhp0857 | *hopK* | 1 | 0 | 0 |
| jhp0870 |  | 55 | 22 | 3 |
| jhp0649 | *homA* | 55 | 18 | 3 |
| jhp0429 | *hopJ* | 2 | 0 | 0 |
| **Genes with alternate stop and start sites** |  |  |  |  |
| **Gene ID** | **Annotation** | **Description** |  |  |
| jhp1031 | jhp1031 | alternate stop, truncated protein in isolate D1 |  |  |
| jhp0440 | jhp0440 | alternate stop, truncated protein in isolate D1 |  |  |
| jhp1103 | *hopQ* | alternate start site, truncated protein in isolate D1 |  |  |

**Table S4.**
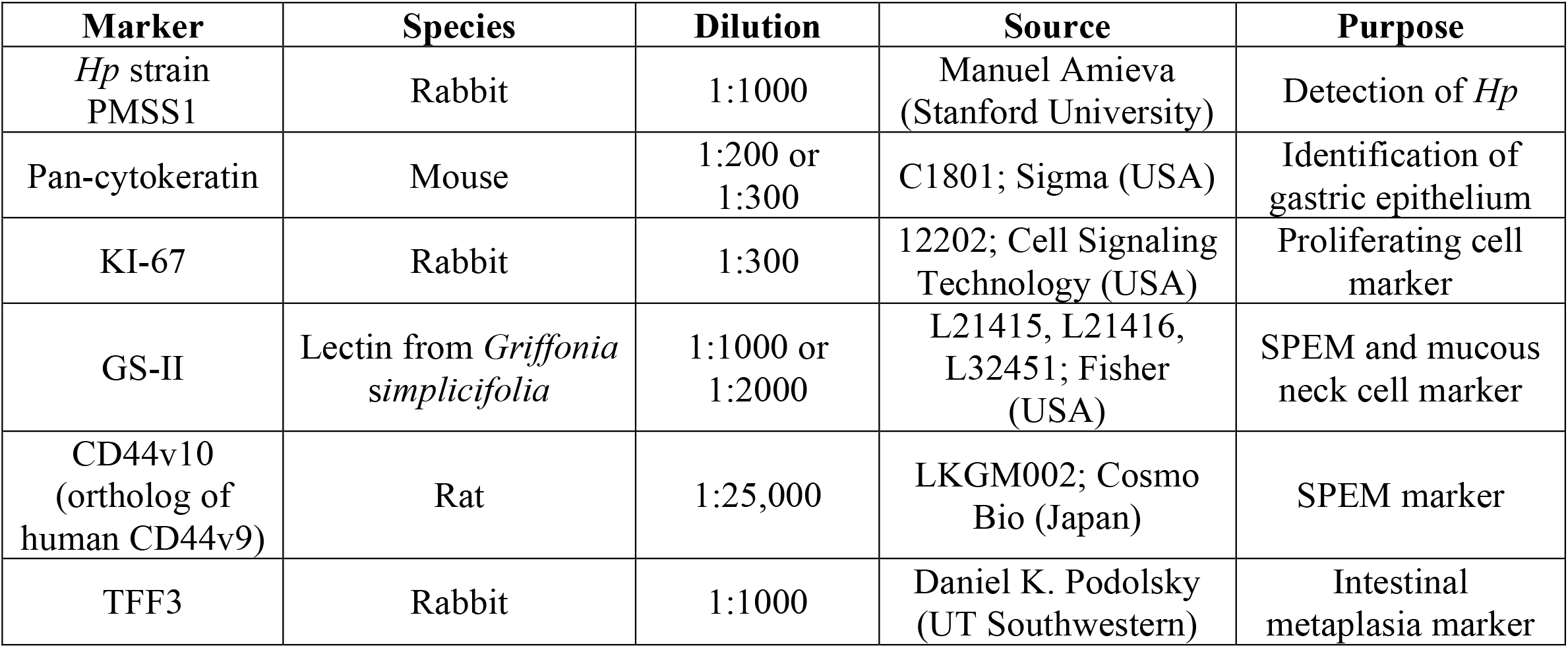
Antibodies and lectins used to assess H. pylori gland colonization and expression of epithelial disease markers.

